# Reference genome-independent taxonomic profiling of microbiomes with mOTUs3

**DOI:** 10.1101/2021.04.20.440600

**Authors:** Hans-Joachim Ruscheweyh, Alessio Milanese, Lucas Paoli, Nicolai Karcher, Quentin Clayssen, Marisa Isabell Metzger, Jakob Wirbel, Peer Bork, Daniel R. Mende, Georg Zeller, Shinichi Sunagawa

**Author notes:** Contributed equally to this work (Hans-Joachim Ruscheweyh, Alessio Milanese).

## Abstract

**Background:** Taxonomic profiling is a fundamental task in microbiome research that aims to detect and quantify the relative abundance of microorganisms in biological samples. Available methods using shotgun metagenomic data generally depend on the availability of sequenced and taxonomically annotated reference genomes. However, the majority of microorganisms have not been cultured yet and lack such reference genomes. Thus, a substantial fraction of microbial community members remains unaccounted for during taxonomic profiling of metagenomes, particularly in samples from underexplored environments. To address this issue, we have developed the mOTU profiler, a tool that enables reference genome-independent species-level profiling of metagenomes. As such, it supports the identification and quantification of both “known” and “unknown” species based on a set of select marker genes.

**Results:** Here, we present mOTUs3, a command line tool that enables the profiling of metagenomes for >33,000 species-level operational taxonomic units. To achieve this, we leveraged the reconstruction and analysis of >600,000 draft genomes, most of which are metagenome assembled genomes (MAGs), from diverse microbiomes, including soil, freshwater systems, and the gastrointestinal tract of ruminants and other animals, which we found to be greatly underrepresented by reference genomes. Overall, two-thirds of all species-level taxa lacked a reference genome. The cumulative relative abundance of these newly included taxa was low in well-studied microbiomes, such as the human body sites (6-11%). By contrast, they accounted for substantial proportions (ocean, freshwater, soil: 43-63%) or even the vast majority (pig, fish, cattle: 60-80%) of the relative abundance across diverse non-human-associated microbiomes. Using community-developed benchmarks and datasets, we found mOTUs3 to be more accurate than other methods and to be more congruent with 16S rRNA gene-based methods for taxonomic profiling. Furthermore, we demonstrate that mOTUs3 greatly increases the resolution of well-known microbial groups into species-level taxa and helps identify new differentially abundant taxa in comparative metagenomic studies.

**Conclusions:** We developed mOTUs3 to enable accurate species-level profiling of metagenomes. Compared to other methods, it provides a more comprehensive view of prokaryotic community diversity, in particular for currently underexplored microbiomes. To facilitate comparative analyses by the research community, it is released with >11,000 precomputed profiles for publicly available metagenomes and is freely available at: https://github.com/motu-tool/mOTUs.

## Background

Identifying and quantifying the abundance of taxa (i.e., taxonomic profiling) is a critical step in linking the composition of microbial communities to environmental functions and host health-related phenotypes [1,2]. Metagenomic sequencing of DNA directly extracted from an environmental or host-derived sample has enabled researchers to taxonomically profile microbial communities in an unbiased and cultivation-independent manner. The development of tools to generate accurate taxonomic profiles from metagenomic data has therefore become important to our understanding of microbial communities [3]. However, existing tools rely on the availability of informative sequences (such as k-mers or marker genes [4,5]), which are predominantly extracted from taxonomically annotated reference genomes (RefGs).

In recent years, high-throughput culturing of microorganisms coupled with RefG sequencing (known as culturomics) [6] has substantially expanded the proportion of microbial taxa with whole genome sequences in data repositories (e.g., NCBI RefSeq) benefitting taxonomic profiling tools. However, there is a strong bias toward microorganisms from well-studied habitats (e.g., human body sites) and/or those that can be readily cultivated using standard laboratory methods. Thus, most microbes on Earth remain uncultivated and lack a representative RefG [7,8], although they can be both globally prevalent [9] and numerically dominant in many environments [10, 11, 12, 13]. As a result, the incorporation of RefGs from newly isolated microbes into taxonomic profiling tools can be slow and disproportional across environments. This poses an additional challenge for accurate taxonomic profiling, given that microorganisms that remain undetected bias the abundance estimates of those that are detected [14,15].

To close the gap between the detectable and actual diversity present in microbial community samples, we developed mOTUs [14,16], a software tool that uses universal, protein-coding, single-copy phylogenetic marker gene (MG) sequences to quantify the taxonomic composition of microbial communities from metagenomic sequence data (for further applications, see also Ruscheweyh et al. 2021 [17]). As these MGs are present in all organisms, they can be identified not only in RefGs, but also in metagenomic assemblies. Conceptually, mOTUs is based on clustering sets of MGs representing individual organisms by sequence similarity into species-level units. In the absence of a generalizable species concept for prokaryotes [18,19], we refer to these units as MG-based operational taxonomic units (abbreviated as ‘mOTUs’).

As an alternative to RefG sequencing, draft genomes are increasingly reconstructed by computational binning of metagenomic assemblies into metagenome-assembled genomes (MAGs [20]) or by sequencing amplified DNA from individual cells, resulting in single cell genomes (SAGs [21]). These cultivation-independent methods have provided genomic access to microbial diversity in previously underexplored environments. Here, in addition to MGs found in RefG and metagenomic data, we now incorporate those found in MAGs and SAGs to more than double the number of taxa represented, adding >20,000 new mOTUs compared to the previous major release [14]. Our evaluations show that mOTUs3 outperforms other methods as assessed using metrics for taxonomic tool benchmarking developed independently from our study [3,22]. Furthermore, we found mOTUs3 to provide an unprecedented view of the species-level diversity within the most dominant heterotrophic bacterial clade in the ocean and to greatly extend the number of detected and differentially abundant species in cross-sectional studies, as exemplified in a comparison between rumen microbiomes of high- and low-level methane-emitting sheep.

## Results

### Taxonomic profiling of diverse environments with mOTUs3

We developed mOTUs3 to facilitate the metagenomic profiling of 33,570 mOTUs, which is a 4.3-fold increase compared to mOTUs2 (Figure 1a). Among all mOTUs, 35% were represented by a RefG (n=11,915; ref-mOTUs), while an additional 21,655 were derived using MGs from either metagenomic contigs (n=2,297; meta-mOTUs) or extended sources, such as MAGs (*de novo*-assembled or imported) and a smaller number of SAGs and isolate genomes (n=19,358; ext-mOTUs), to substantially extend the database coverage for reference genome-independent taxonomic profiling of diverse environments. MGs not assigned to any mOTU were additionally added to the database and merged into a single ‘unassigned’ group to improve the quantification accuracy of taxonomic profiles, as previously demonstrated [14].

**Figure 1.**
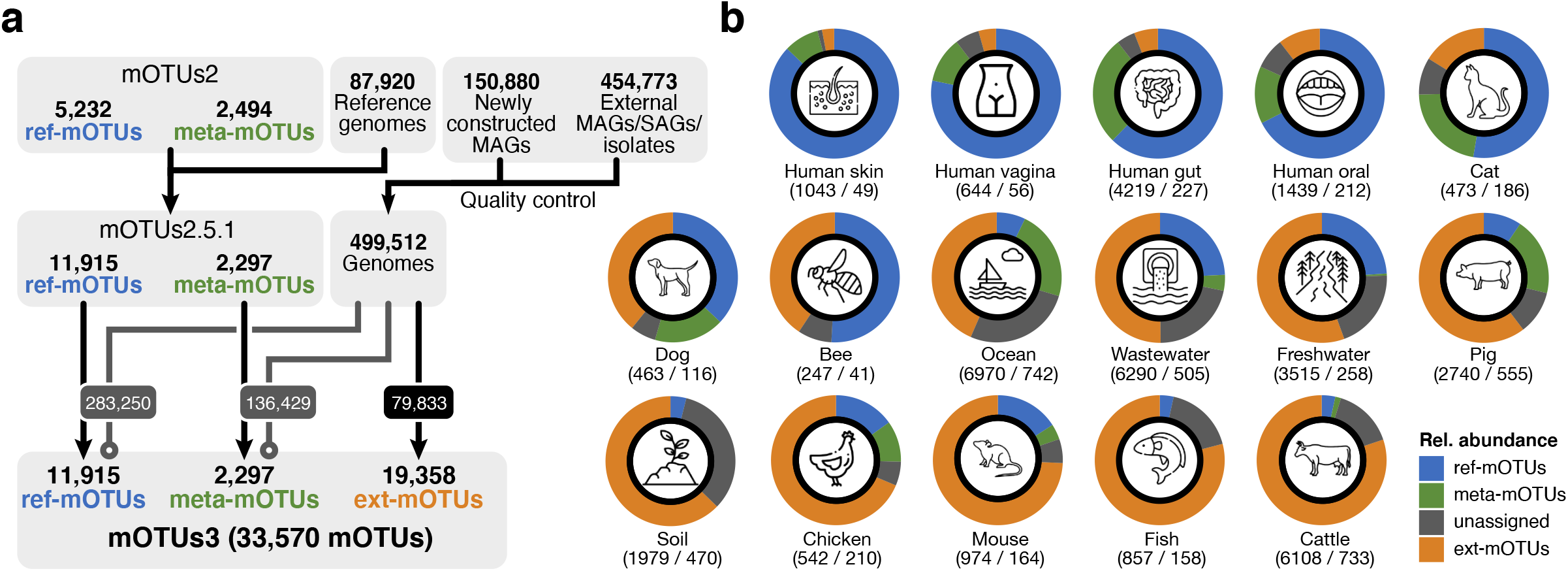
The mOTUs3 database enables species-level profiling across diverse environments. **(a)** The database of the previous major release of mOTUs (version 2)[14] was updated to version 2.5 to account for the current release of the progenomes2 database[49]. Based on version 2.5, the mOTUs3 database was constructed by adding universal, single-copy phylogenetic marker genes (MGs) from 605,653 genomes (metagenome-assembled genomes (MAGs) and a smaller number of isolate and single amplified genomes (SAGs)). This addition resulted in the extension of the database by 19,358 new species-level, MG-based operational taxonomic units (ext-mOTUs). Genomes already represented by ref- and meta-mOTUs in version 2.5 were not added (gray lines). **(b)** Breakdown by the three types of mOTUs shows that mOTUs3 enables the reference genome-independent profiling of a substantial fraction of microbial diversity across different environments. The numbers below the ring charts represent the total number of mOTUs that were detected per environment (left) considering only species with a prevalence of 0.1% and the median number of mOTUs per sample that were detected after downsampling to 5,000 inserts (right).

The newly established database allowed us to determine and systematically compare the fraction of taxa currently not represented by RefGs in various environments. These environments include extensively studied human-associated ones, for which metagenomic studies are complemented by several culturomics efforts (e.g., Lagier et al. [23]). Furthermore, we included data from >20 environmental and animal-associated microbiomes (Supplementary Tables 1 and 2) that have been primarily studied by metagenomic approaches. Overall, we found that more than half (11,882) of all meta/ext-mOTUs (i.e., mOTUs not represented by any RefG) could not be assigned to any known family (Supplementary Table 3; Methods), illustrating the taxonomic novelty covered by mOTUs3. The distribution of the newly included data into ref/meta/ext-mOTUs was highly variable across the different environments (Supplementary Figure 1). As expected, 97% of the ∼400,000 MAGs from human microbiome samples (Supplementary Table 1) had already been represented by 2,360 pre-existing (i.e., ref/meta-)mOTUs (Supplementary Table 4). Notably, the remaining 3% represented 2,750 new ext-mOTUs, showing that novel species can still be uncovered by studying underrepresented populations, dietary habits and/or disease states [24,25]. By contrast, we found that only ∼25% of the 6,479 MAGs from mouse gut metagenomes (Supplementary Table 1) corresponded to pre-existing mOTUs (n=72), despite ongoing cultivation efforts [6]; the remaining 75% were grouped into 587 ext-mOTUs (Supplementary Table 4). However, the vast majority of ext-mOTUs (n=16,021) resulted from the inclusion of other animal-associated (e.g., ruminants, fish, chicken, pig, bee, dog, cat) and environmental (e.g., soil, freshwater, wastewater, ocean, air) microbiomes (Supplementary Table 1) for which the generation of representative RefGs is lagging.

We used mOTUs3 to profile 10,541 available shotgun metagenomic data sets across the 23 environments covered by its database (Supplementary Table 1). For comparative analyses, we subset the data to 5,756 high-quality samples (Methods; Supplementary Table 5) from 16 environments and found the overall number of detected mOTUs to range from 247 (honey bee) to >6,000 (ocean, wastewater and cattle microbiomes). To illustrate the proportion of quantifying taxa currently not represented by RefGs (Figure 1b), we summarized the cumulative relative abundances of unassigned taxa and the different types of mOTUs (ref-mOTUs, meta-mOTUs, ext-mOTUs). The fraction of unassigned taxa was highest for soil samples (33%; s.d. 8%), which reflects the high microbial diversity in soil as well as challenges in reconstructing genomes from this environment [26]. By contrast, more than 87% (s.d. 0.7%) of the relative abundance was represented by ref-mOTUs in human skin samples mainly due to the dominance of few taxa with cultivated representatives [27]. Similarly, the fraction of relative abundance assigned to ext-mOTUs varied considerably between environments: on average, only ∼6% of the bacterial abundance in human-associated samples was assigned to newly added taxa, while this fraction was as high as ∼80% in cattle rumen microbiomes.

### Comparison with other taxonomic profilers

As in other fields of bioinformatics, there is broad consensus that the performance of analysis tools needs to be carefully evaluated. However, best practices (e.g., balancing precision and recall, selecting criteria for ‘best’ performance) are often debated [28,29], and in microbiome research, an agreement on some fundamental concepts (e.g., sequence vs. taxonomic abundance, representation of unknown taxa in ground truth data) is still lacking [30,31]. In an attempt to address some of these issues in a community-driven effort, modeled after successful examples in other fields [32,33], the Critical Assessment of Metagenome Interpretation (CAMI) has provided curated ground truth datasets along with a tool (OPAL) to reproducibly evaluate metagenomic analysis tools [3,22].

Using the latest CAMI datasets with disclosed results [34], we compared mOTUs3 to its prior major release version (mOTUs2) [14] and other selected metagenomic profiling tools (MetaPhlAn3 [5] and Bracken [4,35], Methods) representing conceptually different, well-performing approaches to taxonomic profiling [30]. Using the OPAL tool for scoring and evaluation, we first evaluated presence/absence (F_1_-score) and relative abundance predictions (L1 norm error) at the species level. For the different datasets, which represented samples from five human body sites and the mouse gut microbiome, mOTUs3 and MetaPhlAn3 performed generally better than Bracken and mOTUs2 (Figure 2a/b). At higher taxonomic ranks, mOTUs3 had similar or higher scores than the other tools. For some datasets, taxonomic ranks and tools, there was little to no room for improvements of the F_1_-score or L1 norm error. This may be due to the simulated datasets being mainly based on taxa for which RefGs are available and/or result from incongruencies of taxonomic annotations used by the different profilers compared to the ground truth. In addition to the L1 norm error, OPAL computes additional metrics for profiling quality (completeness, purity, weighted UniFrac error) and summarizes them across taxonomic ranks into a composite score. Based on this evaluation criterion, mOTUs3 outperformed the other tools (Figure 2c), as well as additional tools assessed in the CAMI challenge (Methods; Supplementary Figure 2).

**Figure 2.**
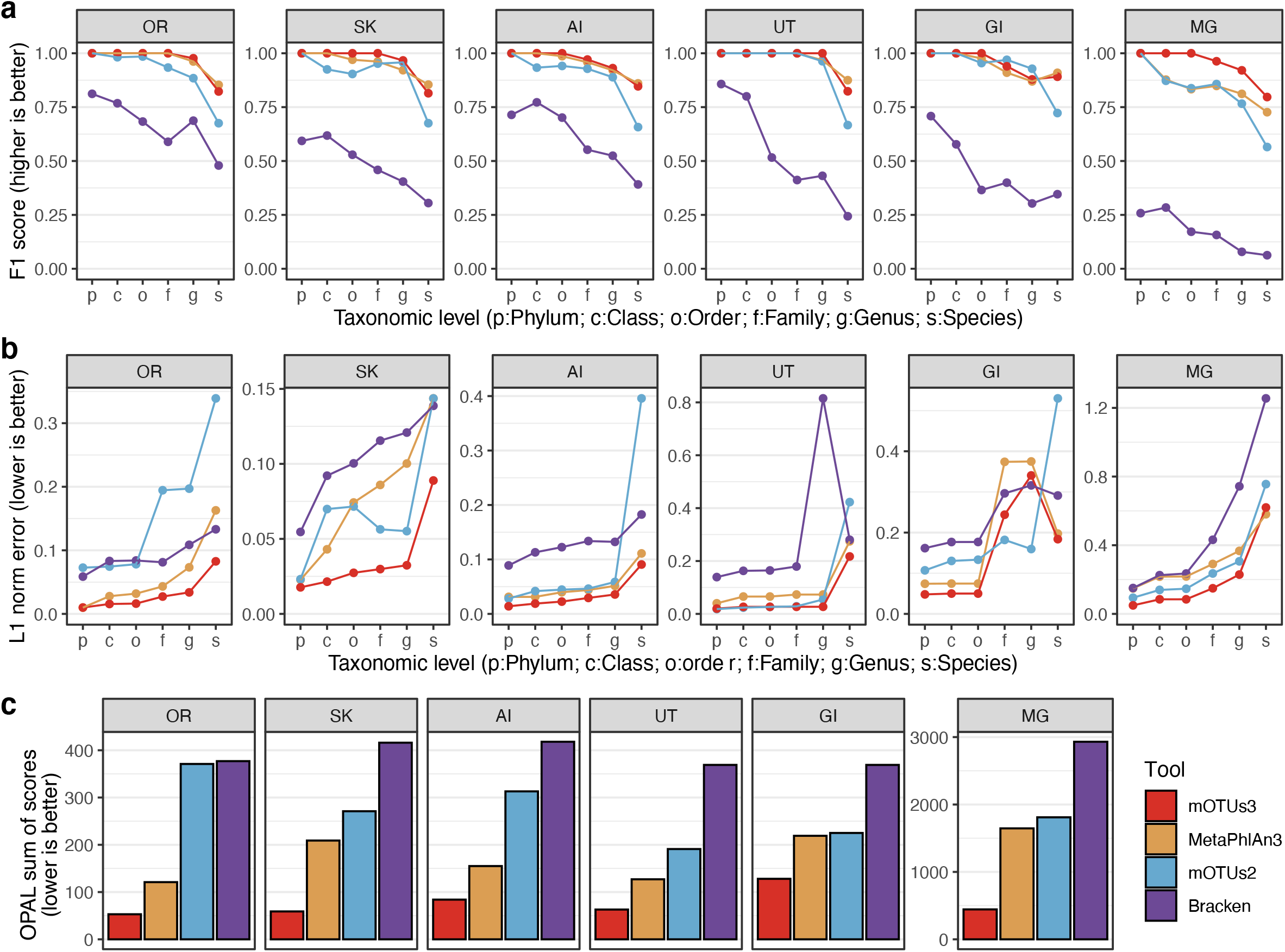
Comparison of mOTUs to other taxonomic profilers. The performance of mOTUs3 was compared to other taxonomic profiling tools based on the dataset from the second Critical Assessment of Metagenome Interpretation (CAMI) challenge (see Methods). The F1 score **(a)** and L1 norm error **(b)** are shown as reported by the OPAL tool[22] for each taxonomic rank (x-axis). High L1 norm error values at the family and genus levels of GI samples mostly derive from an updated taxonomy of the highly abundant Oscillospiraceae (previously Ruminococcaceae)[59]. **(c)** Each method was ranked across all samples and for each taxonomic rank using four measures (completeness, purity, L1 norm error and weighted UniFrac error), and the OPAL sum of scores was calculated as a sum of these ranks (lower rank indicates better performance). OR: oral cavity, SK: skin, AI: airways, UT: urogenital tract, GI: gastrointestinal tract, MG: mouse gut.

In the absence of independent ground truth data sets to benchmark taxonomic profiling tools for less well-studied environments, we correlated taxonomic profiles obtained by mOTUs3 and other tools to those obtained by analyzing 16S rRNA gene (16S) fragments. This approach leverages both the availability of comprehensive 16S databases for taxonomic classification [36] and the possibility of estimating taxonomic abundances based on 16S-based data from metagenomes [37]. Briefly, we extracted 16S fragments from the same datasets we used for metagenomic profiling and generated relative abundance profiles for them (Methods). To ensure comparability between 16S and metagenomic profiles, the analysis was performed at the genus and higher taxonomic ranks (for discussion, see Salazar et al. [37]). We found that mOTUs3 had consistently higher correlations with 16S profiles than the other tools across all environments, except for the human gut for which MetaPhlAn3 showed correlation coefficients similar to those of mOTUs3 (Figure 3).

**Figure 3.**
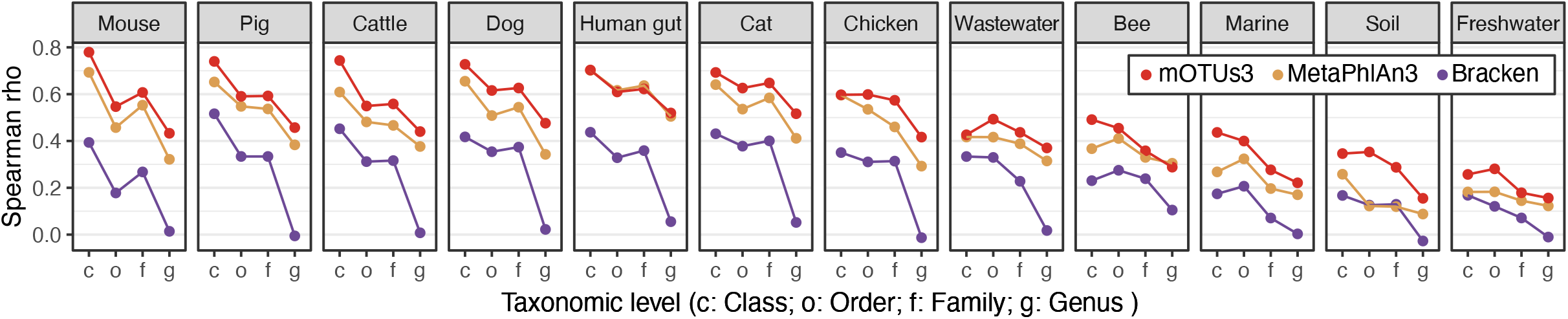
Comparison of metagenomic profiling tools using 16S rRNA-based taxonomic profiles. Spearman correlations between relative abundances generated by different metagenomic profiling tools and 16S rRNA gene-based profiles from the same samples. The correlations were calculated at different taxonomic ranks (x-axis; c: class, o: order, f: family, g: genus) and showed that mOTUs3 generally had the highest values for the different body sites tested, except for human gut samples with similar values for mOTUs3 and MetaPhlAn3.

### Resolving the diversity of Pelagibacterales with mOTUs3

In addition to the broader taxonomic coverage by mOTUs3 across environments, we sought to investigate the capability of mOTUs3 to resolve microbial clades into more fine-grained taxonomic units. To this end, we focused on Pelagibacterales (also referred to as the SAR11 clade), which is the most abundant heterotrophic bacterial group in the global oceans [38]. Members of the Pelagibacterales have previously been shown to display high genomic variability while maintaining highly conserved 16S sequences [39]. This prompted us to evaluate the species-level resolution of mOTUs3 and to compare the diversity represented by mOTUs to the diversity represented by operational taxonomic units (OTUs) defined by 16S sequence similarity.

For this analysis, we selected from all mOTUs annotated as Pelagibacterales (n=1,029; 2,063 genomes) those that were represented by genomes with complete 16S sequences (n=602; 1,105 genomes). The number of mOTUs was comparable to the number resulting from a 95% average nucleotide identity (ANI)-based clustering of the 1,105 genome sequences into species-level groups (n=700; Figure 4a), which is common practice in the field of microbial phylogenomics [7,40]. Moreover, we found sequence identities of mOTUs-representing MGs to linearly correlate with those of whole genomes across the whole range of observed values (*r*^2^=0.71; Figure 4b). By contrast, 16S sequence-based OTUs using a 97% or 99% sequence similarity cutoff resulted in a 31.7-fold (n=19) or 5.8-fold (n=104) lower number of taxonomic units, respectively, compared to mOTUs (Figure 4a). This discrepancy is also reflected by a weaker correlation (*r*^2^=0.45; Figure 4b) of identities between 16S sequences and corresponding whole genome sequences. The minimum 16S identities were ca. 87% and started saturating at approximately 97% at which point genome identities were still as low as ∼70-80% (Figure 4b). Similar findings were reported previously albeit on smaller datasets [39]. Finally, comparing the grouping of genomes by mOTUs and ANI into species-level clusters, we found almost perfect congruence (Figure 4c, Methods).

**Figure 4.**
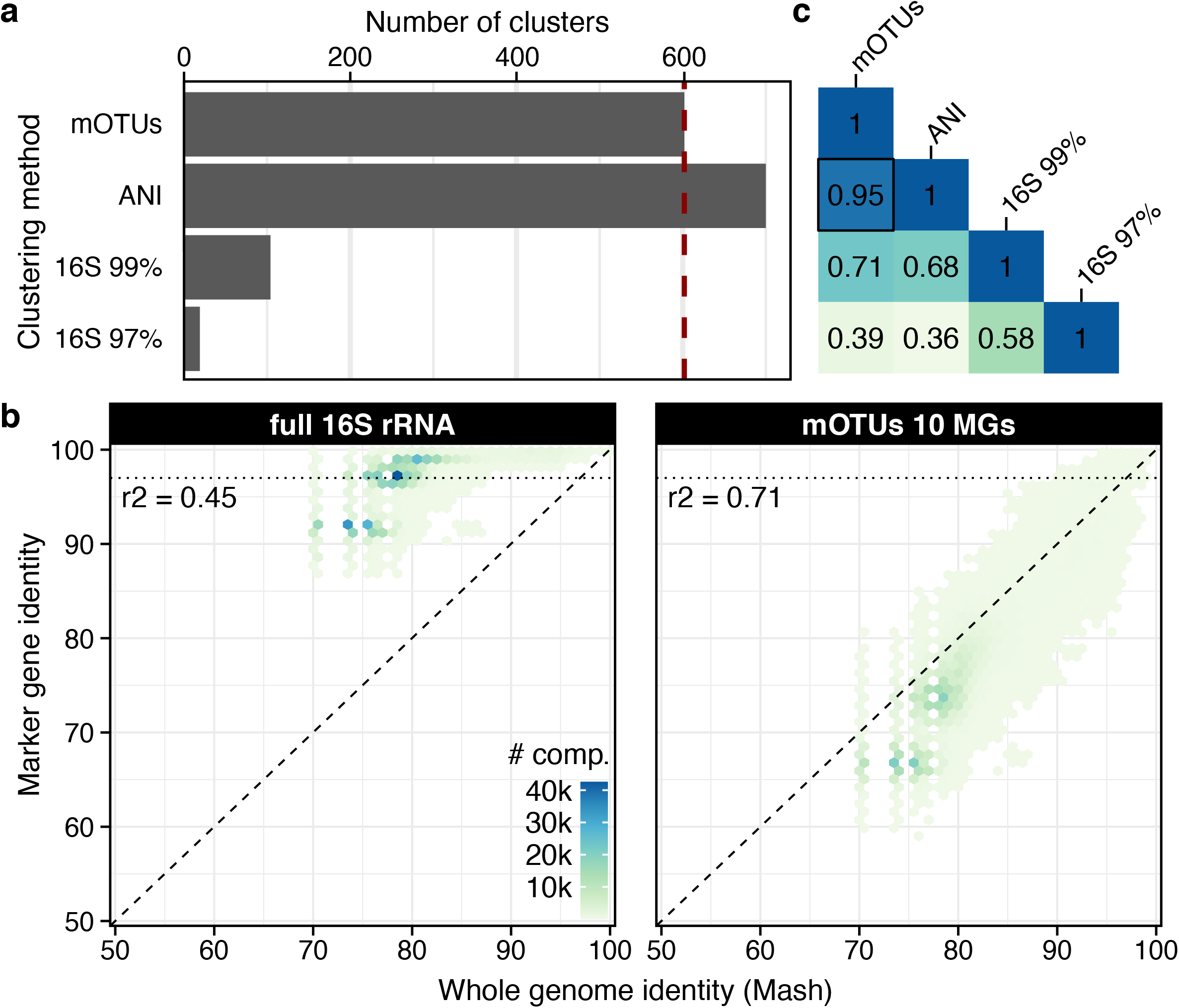
Species-level diversity of Pelagibacterales as resolved by mOTUs3. (a) The number of taxonomic units within the Pelagibacterales order varies depending on the clustering method used, which was based on using marker gene (MG) sequences (used by mOTUs), Average Nucleotide Identity (ANI) of whole genomes, and full length 16S rRNA gene sequences. (b) mOTUs marker gene distances better capture whole genome distances compared to full length 16S, explaining the patterns observed in (a). In particular, 16S rRNA gene sequence identity saturates while whole genome similarity can be as low as 70-80%. (c) The different clustering approaches vary in their agreement with each other as determined by the V-measure, which captures both the completeness and homogeneity of the clusterings. The highest agreement was found between mOTUs and with whole genome clustering by ANI.

### Differential abundance of novel archaea in low/high methane-emitting sheep rumen metagenomes

High-resolution taxonomic profiling of metagenomes from underexplored environments can be achieved by custom-made marker gene or genome databases selected for the microbial community under study [12,41]. However, this approach is often labor- and resource-intensive and requires specialized expertise, and its results cannot easily be compared across studies and communities. To demonstrate the utility of mOTUs3 to address these challenges, we reanalyzed rumen metagenomes from high- and low-methane emitting (HME and LME) sheep [41]. Importantly, these data were not used for the database construction of mOTUs3.

Based on mOTUs3 taxonomic profiles, we identified 131 microbial species that differed significantly in abundance between HME and LME samples and showed an at least tenfold increase or decrease in relative abundance (corresponding to a generalized fold change of >= 1 [42]). Among these differentially abundant species, 92% were represented by ext-mOTUs. These were therefore not expected to be detectable by reference-based profilers. To test this, we applied the same workflow using MetaPhlAn3 and Bracken (see Methods), which yielded only 10 and 30 differentially abundant species for the respective tools (Figure 5a).

**Figure 5.**
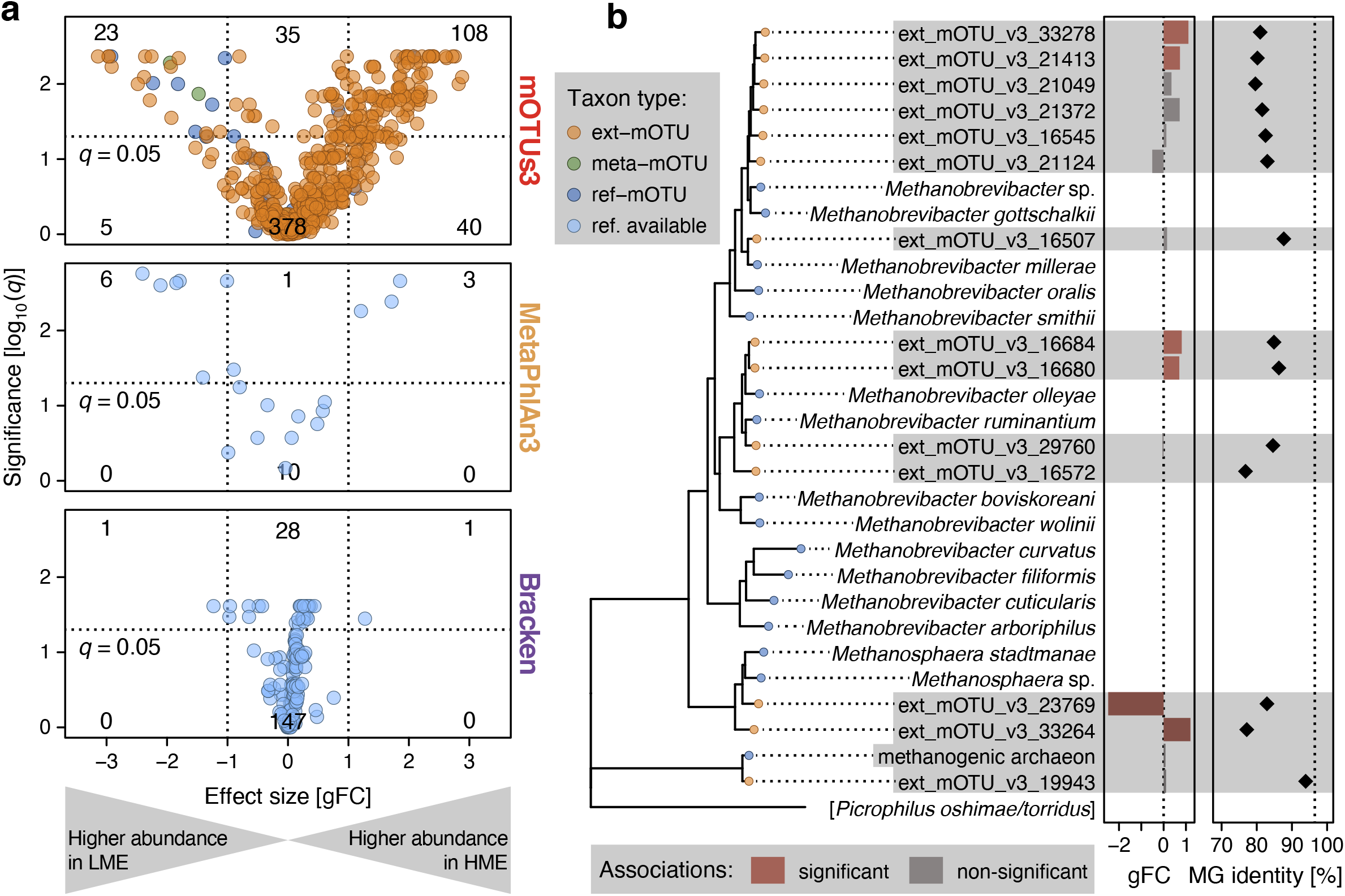
Detection of differentially abundant taxa in low/high-level methane-emiting sheep rumen microbiomes. **(a)** A comparison between metagenomic profilers shows that mOTUs3 detected 131 differentially abundant species (*q*-value <0.05 and an absolute generalized fold change > 1; indicated by dotted lines) between low- and high-level methane-emitting sheep, while MetaPhlAn3 and Bracken detected nine and two species, respectively. Most of the species detected by mOTUs were represented by ext-mOTUs only, demonstrating the added value of reference genome-independent profiling enabled by mOTUs3. **(b)** Archaeal mOTUs present in the sheep rumen microbiome (highlighted in gray) were phylogenetically contextualized with *Methanobrevibacter* spp. and *Methanosphaera* spp. represented by ref-mOTUs. All differentially abundant ext-mOTUs (middle panel) correspond to distinct yet undescribed *Methanobrevibacter* spp. as supported by MG sequence identities (right panel) to the closest known species being below the species-level cutoff of 96.5% (dotted vertical line).

Given the metabolic importance of methanogenic archaea in ruminants as well as previous evidence of uncharted archaeal diversity in the sheep rumen [12], we further investigated the species-level diversity of known and unknown archaeal species. To this end, we reconstructed a phylogenetic tree of the archaeal mOTUs detected in the sheep rumen metagenomes (n=15) and contextualized them with reference genomes from members of the genera *Methanobrevibacter* and *Methanosphaera* (Figure 5b). This analysis revealed that all six differentially abundant archaea in the sheep rumen corresponded to ext-mOTUs. Two of them, which were significantly more abundant in high-methane emitters, were most closely related to *Methanobrevibacter gottschalkii*, which itself was not detected. Notably, the MG sequence similarity between these ext-mOTUs and *M. gottschalkii* was <85% (Figure 5b), which is well below the species-level cutoff of 96.5% used by mOTUs [16] and therefore suggests that these ext-mOTUs represent novel *Methanobrevibacter* spp.

## Discussion

With mOTUs3, we have developed a taxonomic profiler that combines state-of-the-art accuracy, as demonstrated in competitive benchmarks based on simulated datasets, with an innovative database construction approach to detect and quantify underrepresented microbes from diverse environments at high (i.e., species-level) taxonomic resolution. The ability to incorporate MG sequences from any MAG and SAG to generate mOTUs *de novo* and independently from the availability of RefGs and/or prior existence of taxonomic annotations (such as NCBI or GTDB species names) will allow users to continuously extend the core database of mOTUs to represent microbial diversity from newly explored microbiomes. Such future extensions could also target eukaryotic microorganisms, as these are an integral part of many microbial communities, but are not well represented in databases of existing taxonomic profiling tools.

However, the flexibility in defining operational taxonomic units *de novo* comes with a need for taxonomic annotation, as is also the case for 16S rRNA-based *de novo* clustered OTUs. Despite the calibration of MG sequence identity cutoffs to maximize congruence with the NCBI taxonomy [16], this procedure can lead to conflicts with existing taxonomies. Irrespective of the ongoing debate on whether prokaryotic species should be consistent with genomic similarity-based criteria, delineating species by sequence identity puts mOTUs at a disadvantage in benchmarks, such as CAMI, which rely on rigid matching of taxonomic labels. The high performance of mOTUs [34] despite this disadvantage is likely due to the higher number of quantified taxa and the resulting reduction in compositionality-related biases.

## Conclusions

The present work introduces mOTUs3 as a reference-genome independent tool that allows for charting the taxonomic landscape of many environments at species-level resolution. Its independence from taxonomically annotated reference genomes, makes it generally applicable also beyond well-studied environments to quantify and reveal yet uncharacterized microbial species of potential biological relevance. To support the research community, mOTUs3 is documented and available as open source software at https://github.com/motu-tool/mOTUs.

## Methods

### Collection and processing of data to compile the mOTUs3 database

To extend the taxonomic coverage of the mOTUs3 database, 4,531 publicly available metagenomic datasets from 23 environments (Supplementary Table 1) were processed to generate 150,880 MAGs as previously described [43]. Briefly, BBMap (v.38.71) was used to quality control sequencing reads from all samples by removing adapters from the reads, removing reads that mapped to quality control sequences (PhiX genome) and discarding low-quality reads (*trimq=14, maq=20, maxns=1* and *minlength=45*). For metagenomic data of human origin, human genome-derived reads were removed using the masked human reference genome provided by BBMap. Quality-controlled reads were merged using bbmerge.sh with a minimum overlap of 16 bases, resulting in merged, unmerged paired and single reads. The reads were assembled into scaffolded contigs (hereafter scaffolds) using the SPAdes assembler (v3.14 or v3.12) [44] in metagenomic mode. Genes were predicted on length-filtered (≥ 500 bp) scaffolded contigs (hereafter scaffolds) using Prodigal (v2.6.3) [45]. Universal single-copy phylogenetic marker genes (MGs) were extracted using fetchMGs (v1.2; *-m extraction*) [16].

Scaffolds were length-filtered (≥ 1000 bp) and within each study, quality-controlled reads from each sample were mapped against the scaffolds of each sample. Mapping was performed using BWA (v0.7.17-r1188; *-a*) [46]. Alignments were filtered to be at least 45 bp in length, with an identity of ≥ 97% and a coverage of ≥ 80% of the read sequence. The resulting BAM files were processed using the *jgi_summarize_bam_contig_depths* script of MetaBAT2 (v2.12.1) [20] to compute within- and between-sample coverages for each scaffold. The scaffolds were binned by running MetaBAT2 on all samples individually (*--minContig 2000* and *--maxEdges 500* for increased sensitivity). These metagenomic bins were complemented with 454,773 external draft genomes (∼96% MAGs; ∼4% isolate and single-cell genomes) from previous work (Supplementary Table 1). Complete genes in external draft genomes and metagenomic bins were predicted using Prodigal (v2.6.3; *-c -m -g 11 -p single*) and MGs were extracted using fetchMGs (v1.2) *(-m extraction -v -i*).

Metagenomic bins and draft genomes were annotated with Anvio (v5.5.0) [47], quality controlled using the CheckM (v1.0.13) [48] lineage workflow (completeness ≥ 50% and contamination < 10%) and filtered for genomes containing at least six out of the 10 MGs used by mOTUs [16] to produce the dataset of MGs from a total of 499,512 *de novo*-generated MAGs (i.e., quality-controlled metagenomic bins) and external draft genomes used for the construction of the mOTUs3 database.

### Construction of the mOTUs3 database

MGs from 499,512 genomes were mapped against the latest mOTUs database (v2.5.1), which was an update of version 2.0 to account for a more recent release of the progenomes2 database [49] (Figure 1a) using vsearch [50] (v2.14.1; *--usearch_global --strand both --id 0*.*8 --maxaccepts 10000 -- maxrejects 10000*). MGs from a total of 283,250 and 136,429 genomes were assigned to existing ref-mOTUs and meta-mOTUs, respectively. These genomes were removed since they were already represented. The remaining 79,833 genomes resulted in an extension of the mOTUs database by 19,358 new mOTUs (ext-mOTUs). For consistency with the taxonomic annotation of ref-mOTUs, ext-mOTUs were annotated using the STAG classifier (https://github.com/zellerlab/stag, version 0.7; default parameters) trained on genomes in the proGenomes2 database [49] (NCBI taxonomy, version: 8 January 2019). MGs identified on scaffolds that were not binned into MAGs were used to update the ‘unassigned’ mOTU, which contain unbinned MGs that are used to estimate the quantity of unknown species, by aligning these MGs against the extended database using vsearch (v2.14.1; *usearch_global --maxaccepts 1000 --maxrejects 1000 --strand both*). MGs that did not align within MG-specific cutoffs [51] were clustered using vsearch (v2.14.1; *--cluster_fast*) using MG-specific cutoffs and the representative sequence was added to the unassigned mOTU.

### Computation of mOTUs3 profiles for comparative analyses

A total of 11,164 metagenomic and metatranscriptomic samples (Supplementary Table 1, Supplementary Table 2) were quality controlled and merged as described above and profiled with mOTUs3 using default parameters and the *-c* option to build a community resource of taxonomic profiles. For comparative analyses across environments, 5,756 of these samples were used after removing all (n=623) metatranscriptomic samples, metagenomic samples from environments with too few samples (termite, panda, aerosols and bioreactor) or from studies comprising samples from different environments and samples with less than 5,000 mapped inserts. To calculate the total number of detected mOTUs for a given environment, we counted the number of mOTUs with a prevalence greater than 0.1% (Supplementary Table 5). To compare the median number of detected mOTUs across different environments, we downsampled the insert counts to 5,000 using the *rrarefy* function of the vegan package [52].

### Comparison of taxonomic profilers using the CAMI framework

The performance of mOTUs3 was evaluated and compared to mOTUs2 and other taxonomic profilers by analyzing 113 publicly available samples (49 human-associated, 63 mouse gut metagenomes) provided by the second CAMI challenge (https://cami-challenge.org/participate). The samples were profiled with mOTUs3 (v3.0.1; *-C precision*), mOTUs2 (v2.1.1; *-C precision*), MetaPhlAn3 (v3.0.7; *--CAMI_format_output --index mpa_v30_CHOCOPhlAn_201901*) [5] and Kraken/Bracken (v2.1.2; *--db=k2_standard_20201202 --paired /* v2.6.1; *--db=k2_standard_20201202 -r 100 -l S*|*G*|*F*|*O*|*C*|*P*|*D*) [4,35]. Kraken/Bracken reports were further translated into the CAMI format ed files using the *tocami*.*py* script provided at https://github.com/hzi-bifo/cami2_pipelines. For comparative analyses, the OPAL framework (v1.0.9) [22] was used with default parameters providing the gold standard with the parameter *--gold_standard_file*, the names of the tools with *--labels*, the description with *-d*, the output with *--output_dir* and the taxonomic profiles files as positional arguments.

### Comparison of metagenomic profiles with 16S rRNA gene-based profiles

The 16S rRNA-based taxonomic profiler mTAGs [37] (v1.0.1; *-ma 1000 -mr 1000*) was used to generate relative abundance profiles for metagenomic samples (Supplementary Table 1). The output of mTAGs was mapped to the NCBI taxonomy to facilitate comparative analysis. The same samples were profiled with MetaPhlAn3 (v3.0.7; *--index mpa_v30_CHOCOPhlAn_201901*) and Kraken/Bracken (v2.1.2; *--db=k2_standard_20201202 --paired* / v2.6.1; *--db=k2_standard_20201202 -r 100 -l S*). Samples with small read/insert coverages (mTAGs<10,000, mOTUs<1,000, Kraken/Bracken<10,000, no filtering was done on MetaPhlAn3 as profiles contain relative abundances) were removed, leaving 6,119 samples for comparative analysis. Spearman correlations were calculated for each taxonomic rank based on concatenated relative abundances between mTAGs and the metagenomic profiling tools.

### Comparison of Pelagibacterales genome clusters with marker gene and 16S rRNA gene sequences

Out of 2,063 genomes belonging to 1,029 mOTUs annotated as Pelagibacterales, 1,105 genomes (from 602 mOTUs) that contained a complete copy of the 16S rRNA gene were selected. These genomes were also clustered based on average nucleotide identity using dRep [53] (v2.5.4; *-comp 0 - con 1000 -sa 0*.*95 -nc 0*.*2*) using a 95% cutoff as part of the OMD [43]. In addition, these genomes were clustered based on their 16S rRNA gene identity (99% and 97%) using vsearch [50] (v2.14.1; *--cluster_smallmem --id 0*.*97 / 0*.*99*). The consistency between the different clustering approaches was evaluated using the V-measure, which combines both the homogeneity and completeness metrics [54].

To correlate distances of the 1,105 genomes between the different clustering techniques we performed exhaustive distance calculations at the whole-genome level, the 10 MGs used by mOTUs and the 16S rRNA gene. Whole genome distances were computed using MASH [55] as implemented in dRep (v2.5.4). MG- and 16S rRNA gene-based distances were computed using vsearch (v2.14.1; *--allpairs_global --id 0*.*0*) and MG distances were averaged across the 10 genes prior to computing correlations.

### Differential abundance of mOTUs between low/high methane-emitting sheep

Samples from sheep rumen metagenomes (n=16) [41] were profiled with mOTUs3 (v3.0.1; *-c*), MetaPhlAn3 (v3.0.7; *--index mpa_v30_CHOCOPhlAn_201901*) and Kraken/Bracken (v2.1.2; *--db=k2_standard_20201202 --paired* / v2.6.1; *--db=k2_standard_20201202 -r 100 -l S*). To test for differentially abundant species between low methane emitters (LMEs) and high methane emitters (HMEs), the respective profiles were analyzed using SIAMCAT default workflows [42]. This workflow includes filtering of species/mOTUs with a relative abundance of >0.1% in at least one sample [42]. Wilcoxon test results were corrected for multiple testing using the Benjamini–Hochberg method [56] at 5% FDR. The reported effect size measure is the generalized fold change (gFC), calculated as the log10 of the geometric mean of quantile differences between groups as defined in SIAMCAT [42].

A phylogeny was constructed for all archaeal mOTUs belonging to the *Methanobrevibacter* and *Methanosphaera* genera or the *Thermoplasmata* class that passed the relative abundance filtering (14 ext-mOTUs, 1 ref-mOTU) together with ref-mOTUs from *Methanobrevibacter* and *Methanosphaera* (n=15) and a randomly selected Thermoplasmata ref-mOTU as an outgroup. Representative genomes from these 31 mOTUs were selected either by picking the centroid genome (for ext-mOTUs) or the reference genome (for ref-mOTUs). Marker genes were individually aligned (*mafft* [57], v7.458), the alignments were concatenated and a maximum-likelihood phylogeny was calculated using RaxML [58] (v8.2.12; *raxmlHPC -p 12345 -m PROTGAMMAAUTO*). The distance between the 14 ext-mOTUs and their closest ref-mOTU was calculated based on averaged marker gene distances across the 10 genes (v2.14.1; *vsearch --allpairs_global --id 0*.*0*).

## Declarations

### Ethics approval and consent to participate

Not applicable

### Consent for publication

Not applicable

### Availability of data and materials

The mOTUs3 software is documented and publicly available as open source software (GPL 3) at https://github.com/motu-tool/mOTUs. The updated mOTUs3 database can be found at Zenodo (https://doi.org/10.5281/zenodo.5140350) and contains all MGs used in this study and the public profiles generated with mOTUs3. A complete list with all sequencing samples used for building the database and/or for profiling can be found in Supplemental Tables 1 and 2.

### Authors’ contributions

GZ and SS conceived and supervised the work. HJR and AM developed code, generated the database with support from DRM, and performed the benchmark analysis. LP and NK performed the taxonomic diversity analysis of the SAR11 clade and the comparative metagenomic analysis, respectively. QC supported the collection and processing of data. MIM and JW contributed to the taxonomic annotation of mOTUs. HJR, AM, LP, NK, PB, DRM, GZ and SS wrote the manuscript. All authors read and approved the final manuscript.

## Acknowledgments

We would like to thank the ETH IT Services and HPC facilities for granting access to the EULER high performance cluster. We also thank Thea Van Rossum for her input on the taxonomic annotation of mOTUs and the users of mOTUs for their feedback and continuous support.

## Authors’ information

Hans-Joachim Ruscheweyh and Alessio Milanese contributed equally to this work.

## Supplementary Information

Supplementary Information for this manuscript includes:

- Legends for Supplementary Figures 1-2
- Legends for Supplementary Tables 1-5

**Supplementary Figure 1.**
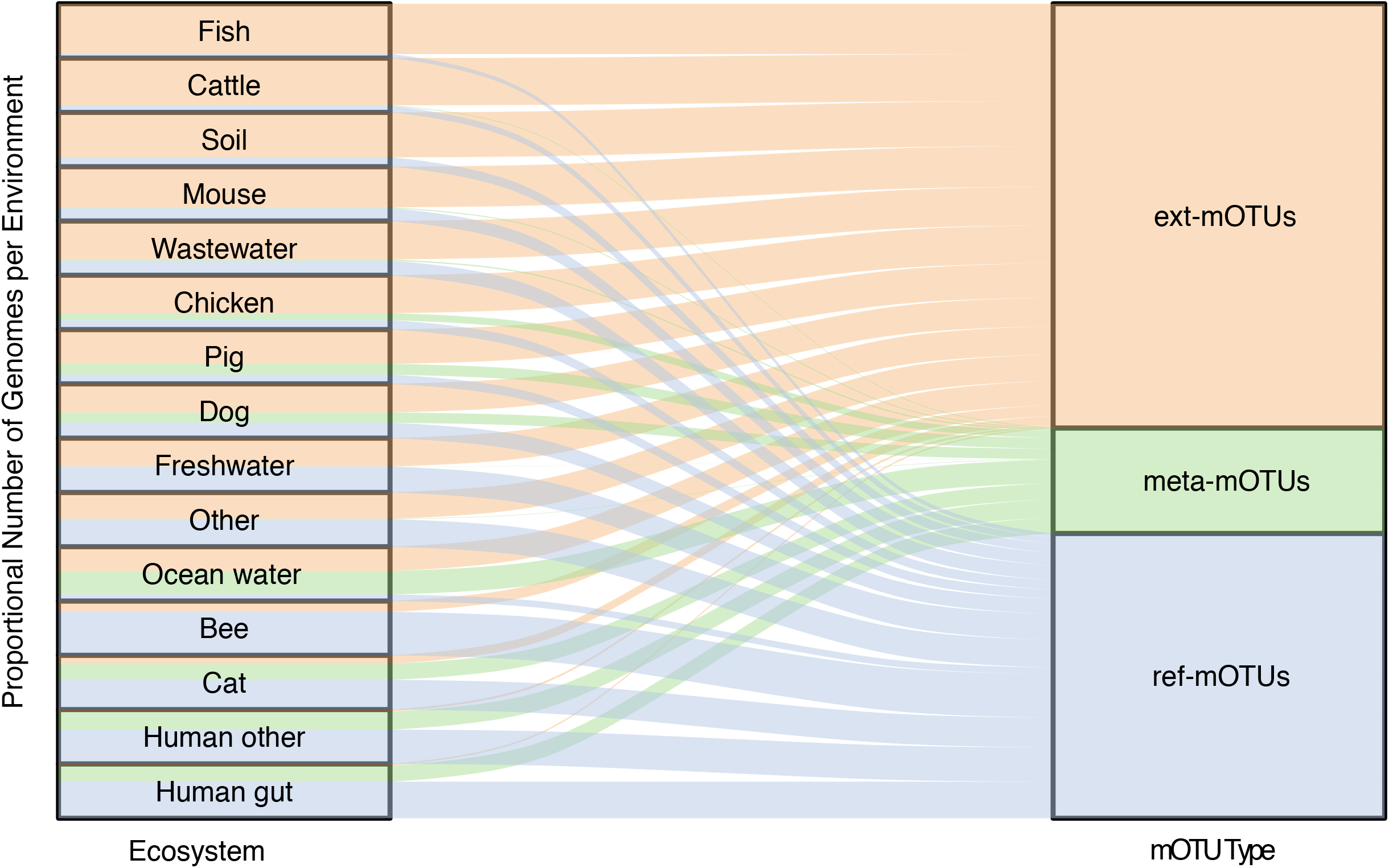
Environment-specific membership of genomes in ref-, meta- and ext-mOTUs. A total of 499,512 genomes derived from 23 environments (environments with few genomes are grouped as ‘Other’, see Supplementary Tables 1 and 3) were used for the extension. The number of genomes was normalized by environments. The proportions of genomes per environment that are either associated with ref- and meta-mOTUs or were used to build ex-mOTUs are shown in the colors blue, green or orange, respectively. For example, the majority of genomes from the human gut match ref-mOTUs, whereas the vast majority of genomes from the fish environment are used to build ext-mOTUs.

**Supplementary Figure 2.**
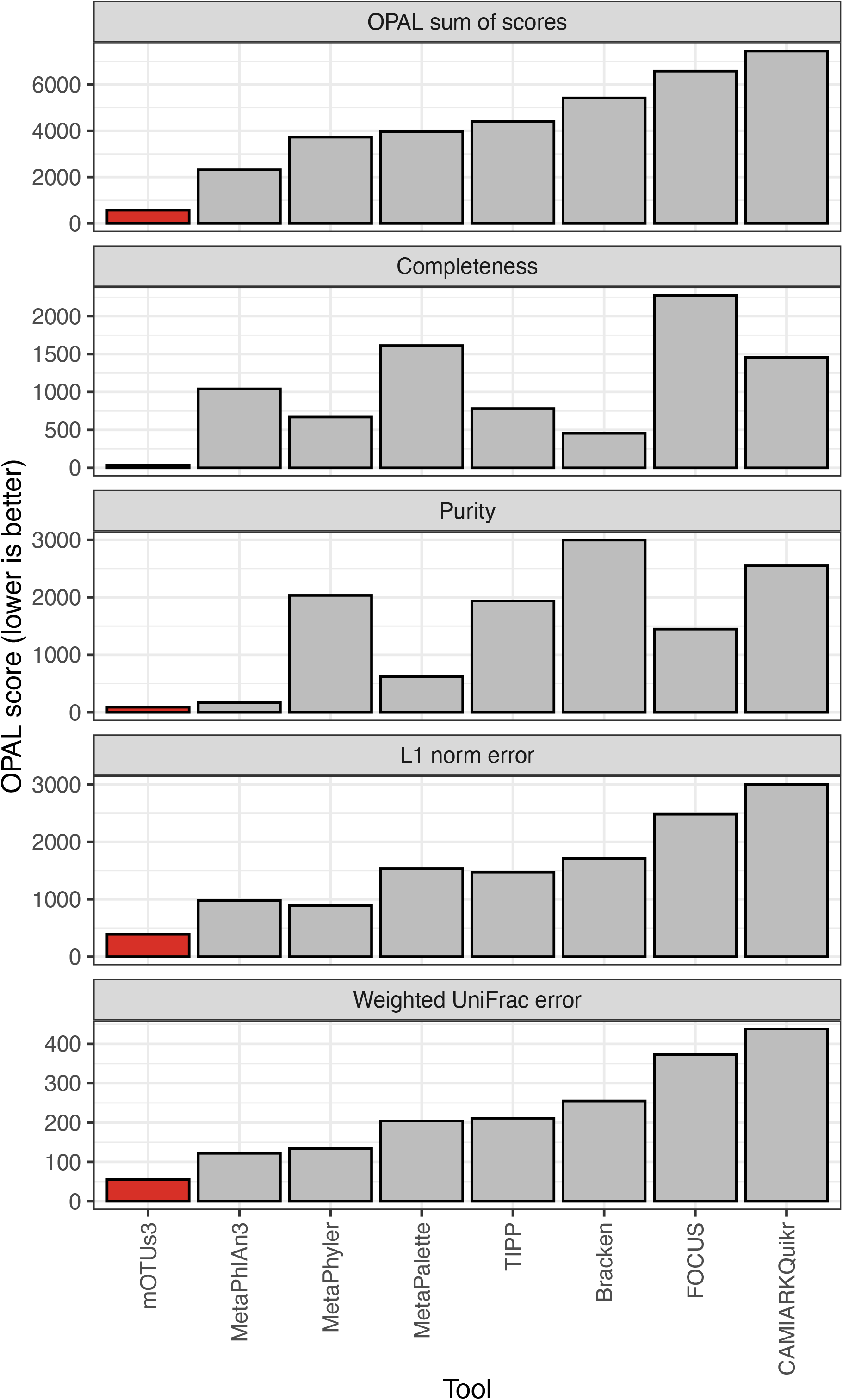
OPAL score broken down to individual metrics for the 63 mouse gut metagenomic samples. The evaluation was performed using the OPAL tool [1] on 63 simulated mouse gut metagenomes [2], which also provided taxonomic profiles for seven different taxonomic profiling tools, and to which we have added mOTUs3 profiling results. The OPAL tool ranks the tools for each sample and for each taxonomic level. The measures considered are completeness, purity, L1 norm error and weighted UniFrac error, shown individually in the bottom 4 plots. Tools with a lower score perform better, as the OPAL score is a sum over rank. The top plot represents the OPAL sum of scores, which is the sum over the four individual measures. mOTUs3 scored best in all categories, including the OPAL sum of scores.

## Supplementary Table Legends

**Supplementary Table 1: Included studies and associated environments**.

Data from 91 studies from 23 environments were included in the extension and/or profiling of the mOTUs database. Of these, 39 studies were selected for in-house MAG reconstruction and 11,164 sequencing samples from 67 studies were used for taxonomic profiling.

**Supplementary Table 2: Sequencing samples included in the taxonomic profile**.

A total of 11,164 samples were taxonomically profiled. Sample names are connected to public repositories by biosample and sequencing run ids. The project name column links the sample name to the study name used in Supplementary Table 1.

**Supplementary Table 3: Breakdown of taxonomic novelty in ext-mOTUs**.

Taxonomic novelty increases with higher ranks, i.e., more than 50% of ext-mOTUs were assigned to previously unknown families.

**Supplementary Table 4: Contribution of genomes to ref-, meta-or ext-mOTUs**.

Genomes/MAGs from different studies and environments contribute in varying proportions to the extension of the database.

**Supplementary Table 5: Data for Figure 1**.

For each sample that passed the filter (total 5,756), we reported the relative abundance for each mOTU type. Additionally, we added the total number of detected mOTUs and the habitat.

